# Increasing Animal Cognition Research in Zoos

**DOI:** 10.1101/2021.11.24.469897

**Authors:** Elias Garcia-Pelegrin, Fay E. Clark, Rachael Miller

**Affiliations:** Department of Psychology, University of Cambridge, Cambridge UK; School of Psychological Science, University of Bristol, Bristol UK

**Keywords:** Animal cognition, animal behavior, environmental enrichment, zoo, welfare

## Abstract

Animal cognition covers various mental processes including perception, learning, decision-making and memory, and animal behavior is often used as a proxy for measuring cognition. Animal cognition/behavior research has multiple benefits; it provides fundamental knowledge on animal biology and evolution but can also have applied conservation and welfare applications. Zoos provide an excellent yet relatively untapped resource for animal cognition research, because they house a wide variety of species - many of which are under threat - and allow close observation and relatively high experimental control compared to the wild. Multi-zoo collaboration leads to increased sample size and species representation, which in turn leads to more robust science. However, there are salient challenges associated with zoo-based cognitive research, which are subject-based (e.g., small sample sizes at single zoos, untrained/unhabituated subjects, site effects) and human-based (e.g., time restrictions, safety concerns, and perceptions of animals interacting with unnatural technology or apparatus). We aim to increase the understanding and subsequent uptake of animal cognition research in zoos, by transparently outlining the main benefits and challenges. Importantly, we use our own research (1) a study on novelty responses in hornbills, and (2) a multi-site collaboration called the ‘ManyBirds’ project to demonstrate how challenges may be overcome. These potential options include using ‘drop and go’ apparatuses that require no training, close human contact or animal separation. This article is aimed at zoo animal care and research staff, as well as external researchers interested in zoo-based studies.

**RESEARCH HIGHLIGHTS:**

- Zoos are an excellent yet relatively untapped resource for animal cognition research.
- Zoo cognition research has historically been challenging, and traditional laboratory paradigms often do not translate well to the majority of zoos.
- Salient challenges of zoo-based cognitive research can be overcome by using less restrictive test apparatuses, limiting animal training and isolation, and subscribing to multi-zoo collaborative programs.

## 1 INTRODUCTION

Animal cognition refers to a suite of mental processes including perception, learning, decision-making and memory (Shettleworth, 2010). To understand cognition, which is essentially ‘invisible’ because it takes place within the brain, we can observe how an animal behaves and make inferences from this. Typically, cognitive research has been undertaken in laboratories because they provide the most highly controlled conditions, using cognitive tasks (i.e., experimental apparatuses). In brief, a cognitive task is designed to permit a restricted number of behavioral responses, and therefore pinpoint whether an animal has a particular cognitive skill. For example, the classic ‘trap-tube’ paradigm consists of a transparent plastic tube containing a food reward and some sort of ‘trap’ through which food can fall, initially developed by primatologists (capuchin monkeys *Cebus apella*, Visalberghi and Limongelli, 1994). We can infer what the animal understands about traps (i.e., a particular aspect of their physical cognition) by how they maneuver food through the tube in relation to the trap. This paradigm has been modified (e.g., ‘two-trap tube’, ‘trap-table’) and tested in a range of species, including primates and birds (e.g., chimpanzees *Pan troglodytes* and children, Horner and Whiten, 2007; rooks *Corvus frugilegus*, Seed et al. 2006; parrots *Psittaciformes*, Liedtke et al., 2011). Overall, there have been multiple critiques of the task apparatus-based approach to studying cognition (as opposed to natural observations; Rowe and Healy, 2014; Heyes, 2015), but it has been the most predominant approach in the literature since the animal cognitive revolution (Shettleworth, 2010). Alternatively, a smaller number of cognitive scientists explore animal cognition in the wild, sometimes using apparatuses, but more often performing observational studies of animal behavior under naturalistic conditions (Byrne and Bates, 2011; Pritchard et al. 2016). This approach has different challenges, such as finding habituated populations to study, quantifying the exact cognitive skills or domains of interest when they are not bounded by a clear task and adapting laboratory tasks for field-use (Pritchard et al., 2016).

Lying at a unique midpoint between laboratory and wild cognitive studies, we find zoos. In zoos, we usually have a high level of animal access (and clarity of observation), and some conditions are easier to bring under control compared to the wild, but not as much as in laboratories (MacDonald & Ritvo, 2016; Hopper, 2017). It is clear from the literature that zoo-based research, including husbandry evaluation, animal cognition, biology, and health, is gaining momentum and scientific outputs (Rose et al., 2019). However, it appears that the type of cognitive research undertaken is strongly laboratory-themed (e.g., high training requirements, close human-animal interactions, heavy reliance on computer touchscreens, Egelkamp & Ross, 2019). It can therefore largely only be undertaken in a handful of dedicated and resource-rich zoos, such as those with permanent research staff or those with direct University links. For example, primate research at the Royal Zoological Society of Scotland and University of St Andrews Living Links/ Budongo Research Consortium in the UK (MacDonald & Whiten, 2011) and the Max Planck Institute for Evolutionary Anthropology and Leipzig Zoo Wolfgang Koehler Primate Research Center in Germany. Interestingly, this may serve to widen the gap between zoos who feel they are capable of cognitive research, and zoos who think they are not.

The overarching aim of this review is to help to change the mindset of zoo professionals and external researchers who may perceive cognitive and behavioral research to be intensive, inaccessible and of little value (such as being “only blue-sky science”, personal communication). We highlight the benefits of cognitive and behavioral research, including applications to animal welfare and conservation research and legislation, animal enrichment, public perception, and education. We will then give a transparent, honest account of the potential challenges, before we demonstrate strategies to address them. The authors are experienced in animal cognition and behavior research, conducted in zoos (e.g., Clark, 2017; Clark et al., 2019; Miller et al., 2014; Miller, Garcia-Pelegrin & Danby, 2021), laboratories and the field. In addition, all authors have worked within UK zoos in other capacities, including as Animal Keepers (R.M. and E.G.P.), as Avian Research Coordinator (R.M.) and Research Officer (F.C.).

## 2 CURRENT BENEFITS AND CHALLENGES OF COGNITIVE RESEARCH IN ZOOS

### 2.1 BENEFITS

The benefits of zoo-based cognitive research center around taxonomic diversity and experimental control. A typical zoo houses several hundred different species, thus providing the highest taxonomic diversity of living specimens of any captive setting. For instance, there is an average of 168 species in a medium-sized Association of Zoos and Aquariums credited zoo (aza.org), and around 400 species are under population management programs within European Association of Zoos and Aquaria zoos (eaza.net). Contrast this to farms and laboratories which typically only house a handful of species or breeds (but with the obvious trade-off of increased sample sizes, Section 3). A recent review of avian cognition research (>500 articles across 30 journals from 2015-2020) indicated that only ~1.4% of bird species were represented, typically from four main orders (Lambert et al., 2021). It thus follows that only 3.9% of these studies were conducted in zoos (74.6% in labs; 17.4% at field sites; 3% at farms and 1.1% did not report the site or were at a mixture of sites; Lambert et al., 2021). Similarly, a review of primate cognition research across a comparable timeframe indicated that only ~13.6% of species were represented (ManyPrimates et al., 2019).

The majority of ‘exotic’ (i.e., non-domesticated or managed animal) cognitive research has taken place on corvids (members of crow family), macaques, great apes, and dolphins (Emery and Clayton, 2004; Tomasello and Call, 1997; Harley et al., 2010), all of which can be found in zoos worldwide but alongside myriad of other taxonomic options. Under most circumstances, it will be easier to access animals living in zoos compared with the wild; particularly rare, dangerous, or cryptic species. Comparing levels of experimental control across different captive settings is less straightforward. In our experience, experimental control is site-dependent, ranging from very high to very low levels of apparatus provision and training within enclosures. There is also the question of controlling non-experimental variables such as climate and human presence. Again, while this depends greatly on the zoo’s ethos and enclosure design, it is probably easier to deal with harsh weather conditions and confounding threats such as predators in a zoo setting compared with the wild.

In terms of cognitive research outcomes, these can broadly be divided into theoretical (or ‘pure’) and applied. Cognitive studies on zoo-housed great apes have made significant contributions to our understanding of human evolution (e.g., Tomasello and Call, 1997), while studies on non-primate cognition have offered important comparative perspectives (e.g., convergent evolution of corvids and apes in Emery and Clayton, 2004). Then, the applications of zoo cognition research can further be divided into animal welfare and conservation. New knowledge of animal cognition has been integrated into animal protection policies. For example, research indicating that fish feel pain has led to increased protection in the fishing and farming industries (Braithwaite, 2010). Similarly, research has led to changes in animal welfare legislation, such as the UK recently extending the scope of Animal Welfare (Sentience) Bill to recognize cephalopod mollusks and decapod crustaceans as sentient beings following an extensive review of the scientific evidence of sentience in these invertebrate animals (Birch et al., 2021; Schnell et al., 2021). Many of the studies with fish and cephalopods were conducted in laboratory settings, however, there is immense scope for other species located in zoos to be studied and hence contribute to more wide-scale protections.

The field of ‘cognitive enrichment’ (Clark, 2011) was barely existent in the literature prior to 2010 but has steadily grown over the past decade within zoos. Cognitive enrichment aims to take knowledge of an animal’s evolved cognitive skills and develop challenges to specifically target these skills. Importantly, animals’ participation in these tasks has been demonstrated to enhance psychological well-being. There are three overarching connections between cognitive challenge and welfare: animals will often seek new challenges and ‘work’ on a challenge without food reward, and solving a challenge is associated with positive emotions or physiological indicators of well-being (reviewed by Clark, 2011, 2017). The results of cognitive research can either feed forward into enrichment design, or research can be designed to simultaneously assess cognitive skill and welfare (Clark et al., 2019). Combining research fields in this way is appealing because it acknowledges the inherent connections between cognition and mental state (Duncan & Petherick, 1991; Mendl & Paul, 2004; Boissy et al., 2007), and maximizes the data collected per subject.

The link between cognition and conservation is also a growing field (Greggor et al., 2014); knowledge of animal cognitive skills and sensory perception has been used to design human-wildlife conflict mitigation strategies in the wild, and to prepare captive animals for reintroduction (Griffin et al., 2000; Maloney and McLean, 1995). Zoos provide access to threatened species that may be otherwise unavailable for cognitive/ behavioral research, which can then be implemented in conservation actions. For example, testing conservation-relevant cognitive abilities, like neophobia (responses to novelty) and innovation (problem-solving) in zoo-housed critically endangered Bali myna *Leucopsar rothschildi*, then implementing these findings in active reintroduction efforts in Bali (Miller, Garcia-Pelegrin and Danby, 2021). In this way, zoo animals have significant opportunities to assist the survival of their wild counterparts.

Finally, other benefits of zoo-based cognition research are best categorized as human-based. Undertaking any research in zoos, cognitive or otherwise, satisfies one of the most important roles of the modern zoo (eaza.net, aza.org; Rose et al., 2019), and feeds forward to educate the public. For instance, the use of cognitive research with dolphins for enrichment, science, education, and conservation at Disney’s The Seas (Harley et al., 2010). More specifically, cognition demonstrating the mental capacities of animals to visitors can raise empathy and be leveraged to discuss wider conservation or welfare issues (Ormandy et al., 2014; Waller et al., 2021). Additionally, the typical approach from researchers is to utilize zoos to conduct research (either themselves or through student projects). We highlight that there are also opportunities for zoo staff to become more closely involved in research, such as data collection, gaining more insight into science, making new connections and, where appropriate, authorship on scientific publications. For instance, zookeepers often know the animals under their care best, and so are well-placed to engage the animals in research participation. However, zoo staff tend to have many tasks and other responsibilities within their roles, and therefore participation is only likely if it is supported by the zoo. Next, we address some of the potential challenges that may impact on cognitive research in zoos, whether conducted by researchers, by non-academic staff, or through collaborative scientific frameworks.

### 2.2 CHALLENGES

The main potential challenges of cognitive research in zoos can be split into animal-based (i.e., concerning the animals used) and human-based (i.e., concerning researchers, care staff, zoo visitors or other stakeholders). We summarize the challenges in tabular form (Table 1) to delineate specific concerns that rarely appear in the literature (we cite where possible), but we have gleaned from our respective experiences in working with and for zoos as researchers and in other roles over the past 20 years. In relation to mitigating these issues, we further outline two specific examples for ‘ideal’ design for zoo-research studies in Section 3.

**Table 1.** Potential Challenges of Cognitive Research in Zoos

| Challenge | Potential Negative Outcomes | Options to Limit/Overcome Challenge |
| --- | --- | --- |
| *Animal-based* |  |  |
| Small sample sizes | Few individuals per species leading to low statistical power (Rutz & Webster, 2020; Button et al., 2020). | -Perform power calculations (e.g., Faul et al., 2009), and seek advice on experimental design to overcome small samples (Saudargas & Drummer, 1996; Dugard & Todman, 2012).<br>-Work with multiple zoos/sites to increase sample size (Many Primates, 2019; Lambert et al., 2021). |
| Untrained/unhabituated animals | Low participation, high drop-out rate, lengthy habituation period required (Melfi et al., 2020). | -Research what motivates the species.<br>-Recruit a larger sample size than needed (i.e., a contingency sample).<br>-Increase the period of habituation (Melfi et al., 2020).<br>-Design experiment with minimal/no training requirement (e.g., “drop and go” apparatuses). |
| Site effect problem (differences between multiple zoos) | Many confounding factors such as differing housing conditions, prior histories, and climate. | -Use repeated measures design so that each animal acts as its own control (Saudargas & Drummer, 1996).<br>-Compare experimental responses to baseline (e.g., how does animal behave without experimental item present; Miller et al., 2021).<br>-Adopt the STRANGE framework (Rutz & Webster, 2020) where individual differences are reported rather than concealed.<br>-Subscribe to a multi-zoo collaborative program like ManyBirds (Lambert et al., 2021). |
| Animal welfare implications of separation, injury, food deprivation or stressful challenge | Animals experience either acute or chronic pain or suffering (Sherwin et al., 2017). | -Ensure research receives full ethical review and is considered by a behaviorist and vet.<br>-Design experiment with minimal/no animal isolation – apparatus could be provided to a whole social group (Gazes et al., 2013).<br>-Continually supervise trials and set criteria for immediately ceasing research if a welfare concern arises.<br>-Provide multiple copies of apparatus if a dominant animal may monopolize access.<br>-Safety-test apparatus on humans first.<br>-Refer to literature on welfare benefits of brief stressful challenges (Meehan & Mench, 2007). |
|  |  | -Build enrichment assessment into cognitive studies (e.g., collect concurrent welfare data to see if cognitive testing can be classified as cognitive enrichment; Clark et al., 2019). |
| Concern over potential long-lasting research implications | Stress-related responses to new people or experimental equipment. Over-familiarity with new people. | -Give special consideration to animals who must retain levels of neophobia or human wariness, i.e., candidates for reintroduction (Griffin et al., 2000).<br>-Undertake habituation to new people and experimental items (e.g., starting outside enclosures or testing without researchers entering enclosure; Kis et al., 2015).<br>-Retain a distance to the animal that matches the distance of usual care staff.<br>-Adapt protocols flexibly if and when required (e.g., number of trials). |
| *Human-based* |  |  |
| Visitor perception and interaction | Lack of understanding of research focus and output. Interference with research. | -Researchers must be professional at all times and prepared to engage positively with visitors.<br>-Recruit assistants to explain the purpose of research, use signage and social media (Waller et al., 2012).<br>-Consider placing research in protected areas and using a video link or changes in height to allow visitors to spectate at a safe distance (Macdonald & Whiten, 2011).<br>-Note that recent study suggests visitors respond more to the behavior elicited by the enrichment than its appearance (Salas et al., 2021) |
| Reluctance | Past negative experience with researchers or perception of research output. | -Hold an initial ‘think tank’ to gain perspectives in a safe space (Gray et al., 2018).<br>-Build relationships based on mutual respect of opinions and expertise.<br>-Set out a memorandum of agreement covering care staff and researcher roles and responsibilities.<br>-Use a third-party advisor to moderate in case of conflict.<br>-Ensure that research outputs suit all stakeholders, e.g., a mixture of peer-reviewed articles, technical papers, and presentations. |
| Time/energy | Research will be another task in an already busy schedule. Where to fit in the research around cleaning, feeding, animal talks, breeding. | -Researchers must be flexible where possible.<br>-Research should require minimal input from care staff (as preferred).<br>-Training should be offered to researchers to move or feed animals (as preferred).<br>-Design research using simple or automated procedures (e.g., remote cameras and hands-free apparatus operation).<br>- Use a drop and go apparatus which can be placed into the enclosure and left with no researcher input. This may require automatically delivering food reward or providing many food containers to access. |
| *Animal- and human-based* |  |  |
| Motivation and Expertise | Subject may have low motivation or a lack of research experience leading to low participation or poor performance.<br>Human may have lack of incentive for research engagement or low familiarity with research procedures and requirements. | -Agree the most suitable animal reward items and frequency of use.<br>-Integrate care staff as active participants in research (where preferred). Provide the appropriate level of co-authorship or acknowledgement (as appropriate).<br>-Zoo management provide time for staff to engage in research as a form of continuing professional development. |
| Dietary | Using food-based rewards for research has likely dietary implications. | -Justify whether food rewards will be additional or part of daily diet.<br>-Time research to coincide with appropriate food delivery times. |
| Breeding or veterinary care | Research could interrupt breeding of threatened species, which is often the priority for many zoo species, or be interrupted if veterinary care/ treatment is required. | -Researchers should obtain necessary permissions (e.g., EAZA ex-situ program coordinators, European studbooks) where applicable prior to data collection<br>-Researchers and zoo personnel may need to co-ordinate around breeding seasons or |
|  |  | veterinary care to reduce any potential disturbance. |
| Health and safety (H & S) | Access to enclosure may be physically unsafe or introduce risk of zoonotic diseases. | -Training on appropriate H & S including safe entry and exit of enclosure, use of safety equipment (where required), regular handwashing, mask-wearing; minimal/ no animal contact.<br>-Provide researchers with radio or emergency number.<br>-Alert auxiliary staff to researcher presence. |

## 3 NOT ALL STUDIES ARE CREATED EQUAL: IDEAL DESIGNS FOR ZOO RESEARCH

Table 1 outlined the main potential challenges and potential options to limit or overcome such challenges when preparing and conducting cognitive research in zoological facilities. Whilst the table outlined does not comprise an exhaustive list of the concerns, it denotes the most likely challenges researchers might encounter when interacting with zoological facilities. Indeed, the researcher endeavoring to conduct cognitive testing with zoological collections ought to understand that contrary to testing within a purposely built laboratory facility, many zoological institutions are less likely to consider research protocols that might be both labor intensive and alter the inner functioning of the animal care team. This poses an initial barrier to the researcher as most protocols in cognitive research entail initial training phases that might take considerable time and may rely on controlling the diet of the subjects under examination. Moreover, facilities in zoo settings are designed with the residing animal and visitor in mind, but not for the occasional cognitive researcher. As denoted in Table 1, a protocol that involves the separation of subjects is unlikely to find success in most zoological settings. Further, small sample sizes may pose an issue as many zoos only have a few exemplars of the desired species due to limited space, species prioritization and other restrictions. Therefore, multi-zoo testing is often a must to get the necessary samples sizes and related statistical power for the study (Button et al., 2013). Multi-zoo research access may require additional permissions. For instance, many zoos belong to professional bodies, such as BIAZA (biaza.org.uk), EAZA (eaza.net), AZA (aza.org) and WAZA (waza.org), that have their own specific protocols and research boards aimed at moderating and facilitating the scientific study of zoological collections.

As we note, there are some challenges of testing in zoological settings that ought to be considered when creating paradigms for zoological testing. The literature already offers prime examples of successful experimental paradigms that address the main challenges one might encounter when researching in zoos. In this section, we will outline two specific examples of the type of research methodologies and frameworks that may offer a powerful set of tools for testing the cognitive capabilities of animals in zoological collections, applicable both to researchers and non-academics interested in engaging in research. We hope that by doing so, we will encourage researchers and zoo personnel to consider zoos and their collections as the great resources for cognitive testing that they can be.

### 3.1 EXAMPLE: CASE STUDY ON NEOPHOBIA IN HORNBILLS

Novelty is the juxtaposition between experience and present stimulus - the more distinct a stimulus is from prior experience, the more novel it will be (Corey, 1978). The discovery of novel items and environments is an unavoidable and important aspect of animal life. This is further enhanced in zoological settings where the ecological and social settings of the animal collections are largely outside of the animals’ control. They may also be liable to change in reference to the zoological facility’s resources, eventualities, and future plans. Animals manage novel input through exploration, which enables the animal to consider the utility of the item. However, novel inputs also pose a potential for danger, as unknown items may not be safe to consume or interact with, and unacquainted species may be predators (Greenberg & Mettke-Hofmann, 2001). Consequently, a degree of caution when deciding to interact with a novel stimulus aids the animal in balancing the potential of its utility against the risk of danger.

Neophobia (from the ancient Greek *Neo* (new) and *phobia* (fear)) refers to the fear that animals present when encountering a novel stimulus. Understanding how animals respond and approach novel stimuli is both vital for cognitive research and conservation because the effects of urbanization will likely force species into new ecosystems where they will have to inherently adapt (Greggor et al., 2014). Moreover, understanding these reactions within zoological collections will also be of use for the animal care team, as it will provide insightful information regarding the degree of mailability in their ecosystem that a species or individual within their collection is likely to endure. For example, an animal with a high level of neophobia towards new foods might require a longer process of food habituation when faced with new changes in their dietary requirements than a species or individual with lower levels. Similarly, a species with high object-level neophobia, such as a member of the corvid family (Miller et al., 2021) may require more habituation steps when changing features in their enclosure than a species with low object neophobia, like a kea *Nestor notabilis*.

In this case study, we investigated the neophobic responses of 9 captive hornbills (*Bucerotidae*, five species) at Birdworld Zoo in Farnham, UK. The zoo granted us access to their hornbill collection for 3 subsequent days, with 20 minutes per day (per enclosure). We selected this case study as an example for several reasons:

#### No need for training

In Table 1, it is highlighted that as zoological collections mainly comprise of untrained and unhabituated animals, this can result in overall low participation rates which can lead to small sample sizes. This neophobia paradigm investigated the natural behaviors of the sample i.e., how quickly they feed on their regular diet in the presence and absence of a novel item, as such, it required no training from the subjects. Moreover, as the paradigm investigates the behavioral reactions of the subjects to a new stimulus, it required no habituation periods, which allowed the experiment to be performed within the time constraints set by the zoological facility.

#### Adaptable for multiple zoo testing

As mentioned in Table 1, one of the main challenges of performing cognitive research in zoological facilities is the small sample sizes in the zoo and the potential issue of site differences (e.g., enclosure size, mixed species exhibits). As this paradigm was quick to perform, simple and did not require any human-subject interaction, it was deemed a perfect methodology for further testing with multiple zoos (e.g., similar approach with Bali myna in Miller, Garcia-Pelegrin and Danby, 2021). Furthermore, one simple way to manage site differences is to compare the experimental responses to a control baseline within each site, which is possible with this paradigm as it includes a familiar food condition (i.e., presented without a novel item).

#### Animal welfare implications

Animal welfare implications should be a priority when designing an experimental paradigm for any facility (be that laboratory or zoological). Zoological collections are often very understandably reluctant to separate their subjects even temporarily (as social animals may not be used to being alone and may react negatively). This neophobia paradigm allowed for social testing as the measurement was a comparison between the responses of the target individual in the control condition with their responses in the experimental condition, both of which were tested within the same pair.

#### Time/energy

Zoological facilities are often busy places that may lack time and staff resources to appoint a dedicated zookeeper to overview the researcher experimenting. Consequently, a successful paradigm for zoological research is likely to be quick to complete and not demand a lot of resources from the zoological team. The paradigm performed in this case study took only 20 minutes per day (per enclosure) over 3 days to complete - under 5 hours in total. Moreover, as the paradigm required the novel food and objects to be placed alongside their daily feed, this was done during their routine feeding time and thus caused no/minimal disruption to the animal care team’s routine.

##### Methods

The neophobia experiment consisted of three conditions that varied depending on the stimulus presented to the subjects. In the control condition, we presented the subjects with their regular diet inside a familiar food bowl. The novel object condition consisted of the addition of a purposely made novel object next to their familiar food. The novel objects were crafted to resemble the items used for neophobia testing in corvids and Bali myna research (Miller et al., 2021; Miller, Garcia-Pelegrin and Danby, 2021), but adapted for the hornbills’ typically larger size (Figure 1A). The novel food condition consisted of the addition of a 10cm^3^ block of orange colored jelly placed inside a secondary food bowl next to the familiar food (Figure 1B). We were interested in the hornbills’ latency to approach and touch the familiar food in reference to the stimulus (or absence of) presented to the subjects. To do so, we measured the behavioral reactions of the hornbills to the novel food and novel object in contrast to when presented with the familiar food alone. We deemed the commencement of each trial as the moment the experimenter left the video shot. Each trial lasted 20 minutes in total (as per Miller, Garcia-Pelegrin and Danby, 2021), which was based on pilot trials where we checked that subjects would approach familiar food alone (no novel items present) within this time frame.

**Figure 1.**
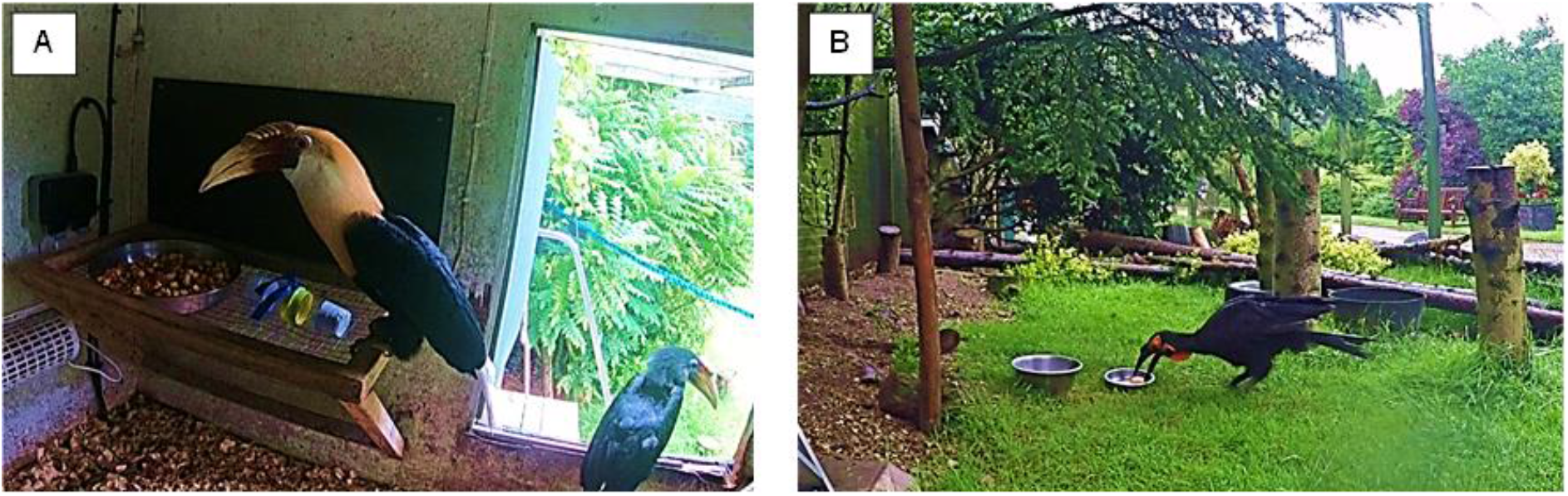
(A) Novel object condition with a pair of Blyth’s hornbills (*Rhyticeros plicatus*). (B) Novel food condition with a Southern ground hornbill (*Bucorvus leadbeateri*) that is pecking at the orange jelly.

##### Subjects

The subjects of this case study were all the hornbill species presently available for testing at Birdworld Zoo (Farnham, UK). The species and sample sizes are displayed in Table 2.

**Table 2.** Hornbill species, sample sizes and housing.

| Species | Binomial nomenclature | Sample size | Housing |
| --- | --- | --- | --- |
| Southern ground hornbill | *Bucorvus leadbeateri* | 2 | Paired |
| Blyth's hornbill | *Rhyticeros plicatus* | 2 | Paired |
| Visayan hornbill | *Penelopides panini* | 2 | Paired |
| Black hornbill | *Anthracoceros malayanus* | 2 | Paired |
| Rhinoceros hornbill | *Buceros rhinoceros* | 1 | Alone |

##### Data Analysis

We video recorded all trials and coded all videos using Solomon Coder (Péter, 2019). To investigate the effects of condition (control, novel food, novel object) on the hornbills’ latency to a) approach and b) touch the familiar food, we conducted a Generalized Linear Mixed Model (GLMM) in RStudio for Mac (version 1.2.1335) with subject as a random effect and condition (control, novel food, novel object) as main effects, using likelihood ratio tests (drop1() function) and Tukey comparisons for post-hoc comparisons (package multcomp, function glht()).

##### Ethics Statement

This non-invasive behavioral bird study was conducted adhering to UK laws and regulations and covered under a University of Cambridge non-regulated procedure. Additionally, we obtained permission to conduct the study from the representatives of the study site (Birdworld, UK).

##### Results

Latency to approach and touch the familiar food differed across the conditions (GLMM: latency to approach, *χ*^2^=7487.1492, df=2, p<0.001: latency to touch *χ*^2^=7572.239, df=2, p<0.001). Hornbills took longer to approach the familiar food when a novel object or novel food was present compared to the control condition (Tukey contrasts: novel object – control, z=78.54, p<0.001; novel food – control, z=−15.03, p<0.001). They also took longer to approach the familiar food when a novel object was present than when a novel food was present (Tukey contrasts: novel object – novel food, z=78.37, p<0.001) (Figure 2).

**Figure 2.**
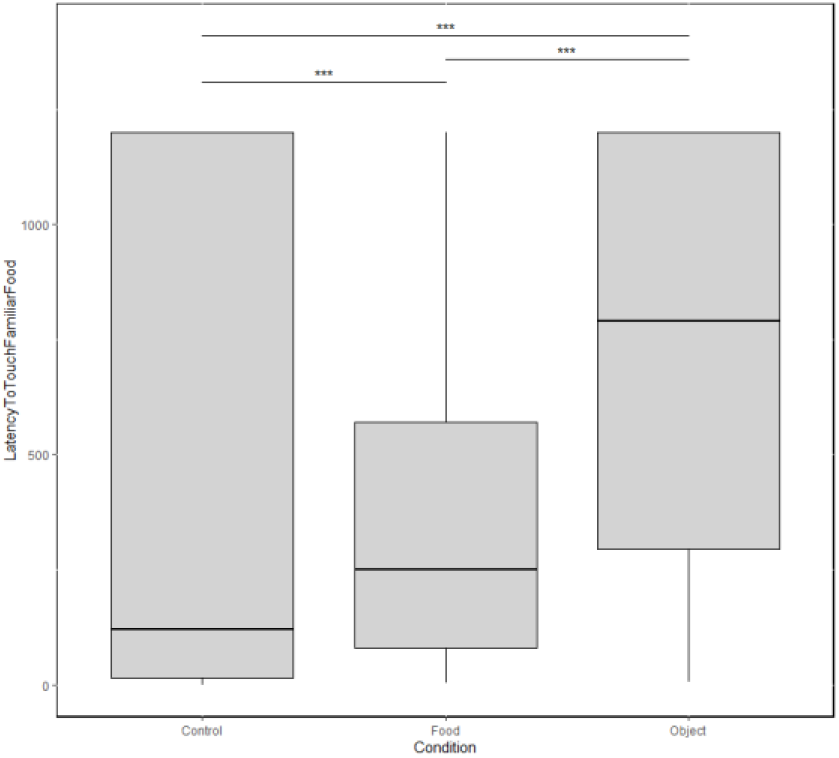
Latency to touch the familiar food differed between conditions. Raw data: lines represent median. *** p < 0.001

##### Discussion

In this case study, we tested the latency to approach and touch familiar food in presence of a novel object or a novel food (i.e.., neophobia) in 5 different species of captive hornbills (9 subjects). We found that hornbills took longer to approach the familiar food when there was a novel object next to it, compared to a novel food, or no novel item present. Additionally, they took longer to approach when a novel food was present than the familiar food alone. These results suggested that hornbills exhibit a degree of caution when faced with novel inputs, which is likely to be a behavioral adaptation that moderates the possibility of danger in the wild (Greenberg & Mettke-Hofmann, 2001; Mettke-Hofmann et al., 2002).

Anecdotally, whilst only 1 subject touched the novel object, 4 of 9 subjects (44.5%) pecked at the novel food (jelly). Using a comparable paradigm, only 20% of (241) corvids (Miller et al, 2021) and 0% of (22) Bali myna touched the novel food (Miller, Garcia-Pelegrin and Danby, 2021). This finding in hornbills may be related to their mainly frugivorous diet (Naniwadekar et al., 2015), which leads them to be more likely to identify the vibrant colors or smell of the orange jelly as a potential food source. The finding that hornbills displayed some neophobic sensitivity when confronted with a novel input should be kept in mind when making changes in their enclosure or diet, suggesting that a period of habituation before any major change is likely a good course of action.

We conducted this study as an example of an experimental design that is relatively simple to perform in zoological settings. We encountered no major technical issues with the protocol. On this occasion, the zoological facility allowed for the experimenter to perform the daily feeds to the birds alongside testing. Being able to present the food to the hornbills (rather than a zookeeper needing to be present to do this) aided in the efficiency of the experiment. We note that the neophobic reactions to new people or keepers presenting food have yet to be studied, though there is some indication that birds respond differently to familiar versus unfamiliar people (Cibulski et al., 2014). This is worth further exploration as it may affect the results of some studies and the willingness of some subjects to participate in the study. Indeed, one of the subjects in the present experiment, the Rhinoceros hornbill, initially displayed some hostile behavior towards the experimenter, though this quickly dissipated in subsequent trials. Thus, it may be worthwhile considering habituating the subjects to a new experimenter if required to enter the enclosure prior to testing. Alternatively, this experiment could be conducted by a familiar zookeeper, as it only required adding a novel stimuli beside their daily feed for a short period of time (20 minutes) across a 3-day period.

Overall, we chose to present this study as an example of the type of simple paradigm that can be informative while mitigating some of the main issues one ought to consider when performing cognitive research with zoological collections (as highlighted in Table 1). While the data alone offers an insightful window into the behavioral reactions of hornbills displayed when presented with certain types of novel stimulus, it is important to note that there are limitations with the data presented. The main limitations of this dataset concern the sample size and lack of repeated testing which makes the results of this study constrained in their inference capability. As denoted in this paper, one way around this issue is through multi-zoo research and/or collaborating with frameworks like the Many X Initiatives (see next section). When doing so it is imperative to consider the inherent differences between the zoos and aim at minimizing them through methodological consistency. For instance, comparing experimental conditions to a control condition (as performed in this case study), enacting a repeated measures design thus treating each subject as their own baseline, or a combination of both.

### 3.2 EXAMPLE: MANY X INITIATIVES

Many X initiatives such as ManyBabies, ManyPrimates, ManyDogs and ManyBirds Projects share a common approach to facilitate large-scale, international collaborative research under Open Science based framework (Lambert et al., 2021; ManyPrimates et al., 2019; Byers-Heinlein et al., 2020). Many of these projects aim to explore the evolution of cognition within specific animal groups, with the potential in future for cross-project collaborations. The Many X projects aim to be inclusive, inviting collaboration between academics and non-academics, across the world with clear, coherent, and accessible frameworks for research participation. The ManyBirds Project, in particular, aims to facilitate collaboration with a variety of potential sites, including zoos, labs, field, and private homes, by selecting experimental designs that are low time and labor intensive, requiring no/ minimal physical contact with the experimenter and therefore suitable for unhabituated/ untrained birds (Lambert et al., 2021). An example of this is the upcoming ManyBirds study on neophobia in birds - following a similar protocol as the above hornbill pilot study (for more information: Twitter: @TheManyBirds; Website: www.themanybirds.com).

## 4 CONCLUSION

Animal cognition and behavior research has important implications for applied sciences including animal welfare and conservation, as well as in education, ethics, and legislation. It can also be enriching for captive animals to participate in research. There is a need to increase species and sample size representation in research (e.g., ManyPrimates et al., 2019; Lambert et al., 2021), and zoological collections provide a unique opportunity to achieve this together in a mutually beneficial manner. We identify some of the relevant benefits and challenges to zoo-based research, and outline potential mitigating options, including specifics for experimental designs that may be most suitable for many zoo environments. We hope that this article contributes to increasing zoo-based cognitive and behavioral research.

## ACKNOWLEDGEMENTS

Many thanks to Birdworld (Farnham, UK) for access to hornbills for this pilot study, particularly to Duncan Bolton, Kat Nicola, and Polly Bramham. Thank you to Gavin Harrison for feedback on a manuscript draft, and Katy Lee Jones for help with the hornbill testing. R.M. and E.G.P. were funded by a Career Support Fund (University of Cambridge) awarded to R.M. At the time of writing, F.E.C was an honorary research associate at the University of Bristol.

## DATA AVAILABILITY STATEMENT

The hornbill study data sheet is available at: https://osf.io/sc684/?view_only=e5897a4277a445a3887bdda6fa3f7d05

